# The magnitude of airway remodelling is not altered by distinct allergic inflammatory responses in BALB/c vs C57BL/6 mice but matrix composition differs

**DOI:** 10.1101/2020.10.20.347096

**Authors:** James E Parkinson, Stella Pearson, Dominik Rückerl, Judith E Allen, Tara E Sutherland

## Abstract

Allergic airway inflammation is heterogenous with variability in immune phenotypes observed across asthmatic patients. Inflammation has been thought to directly contribute to airway remodelling in asthma, but clinical data suggests that neutralising type 2 cytokines does not necessarily alter disease pathogenesis. Here, we utilised C57BL/6 and BALB/c mice to investigate the development of allergic airway inflammation and remodelling. Exposure to an allergen cocktail for up to 8 weeks led to type 2 and type 17 inflammation, characterized by airway eosinophilia and neutrophilia and increased expression of chitinase-like proteins in both C75BL/6 and BALB/c mice. However, BALB/c mice developed much greater inflammatory responses than C57BL/6 mice, effects possibly explained by a failure to induce pathways that regulate and maintain T cell activation in C57BL/6 mice, as shown by whole lung RNA transcript analysis. Allergen administration resulted in a similar degree of airway remodelling between mouse strains but with differences in collagen subtype composition. Increased collagen III was observed around the airways of C57BL/6 but not BALB/c mice while allergen-induced loss of basement membrane collagen IV was only observed in BALB/c mice. This study highlights a model of type 2/type 17 airway inflammation in mice whereby development of airway remodelling can occur in both BALB/c and C57BL/6 mice despite differences in immune response dynamics between strains. Importantly, compositional changes in the ECM between genetic strains of mice may help us better understand the relationships between lung function, remodelling and airway inflammation.

## Introduction

Asthma is a global health problem with increasing prevalence, currently affecting over 300 million people.^1^ Of note, the term “asthma” encompasses a range of disease phenotypes. Progress in understanding heterogeneity of airway inflammation has led to defined asthma endotypes^2,3^ often characterized by the presence or absence of type 2 eosinophilic inflammation and/or type 17 neutrophilic inflammation.^4^ A clarified definition of such inflammatory phenotypes in asthma has led to development of innovative therapies directed at modulating specific inflammatory pathways, in particular type 2 inflammation.^5^ Whilst the use of biologicals targeting IgE or either type 2 cytokines Interleukin (IL)-4, IL-5, IL-13 or their cognate receptors (IL-4Rα, IL-5R, IL-13Rα1) have been effective at reducing disease exacerbations in allergic asthmatics, these therapies are often insufficient to improve underlying disease pathogenesis.^6–8^ Furthermore, clinical trials with antibodies targeting IL-17 signalling have shown no benefit^9^, despite a strong association of severe asthma with type 17 neutrophilic inflammation.^10^ To achieve progress in treating asthma we need a more comprehensive understanding of pathology beyond viewing inflammation as the main instigator of disease.

Along with airway inflammation, asthma is characterized by airway hyper-responsiveness (AHR) and airway remodelling, a process of changes to the composition, content and organisation of cells and extracellular matrix in the lung. Although tissue remodelling is a critical process during development and tissue repair^11^, it is also a pathogenic response in diseases like asthma, and undoubtedly impacts on lung function.^12^ Comprehensive research using a combination of mouse models and human studies typically attribute the development of remodelling to chronic airway inflammation.^13–17^ However, this view conflicts with emerging data that shows remodelling can occur as a primary event prior to inflammation.^18–20^ Glucocorticoid steroids can generally improve lung function but do not alter airway remodelling.^21^ Alternatively, drugs that successfully target specific inflammatory pathways fail to improve lung function^7,8^ presumably because they do not affect airway remodelling. Overall, the links between inflammation, remodelling and lung function are still unclear and thus warrant investigation. Airway remodelling is a complex disease process, difficult to study in patients especially in the context of understanding multiple components that may influence development of individual remodelling processes over time. Therefore, there is a crucial need for animal models that reflect asthma disease processes with a focus on studying the development of stable and irreversible airway remodelling.

Genetic differences between inbred mouse strains are well known to strongly affect both airway inflammation^22–25^ and AHR.^26^ For instance, C57BL/6 mice are known to have a high airway resistance in response to methacholine challenge independent of allergic inflammation^22^, whereas BALB/c mice generally exhibit greater airway reactivity in response to allergens.^26^ Therefore, studies examining airway pathology in different mouse strains can provide a basis to explore the relationships between immune cell dynamics in relation to changes in airway remodelling and lung function. In this study we utilised a model of chronic allergic airway inflammation that shares features of disease common to severe asthma in people, including mixed type 2 and type 17 airway inflammation, steroid resistant neutrophilia and AHR independent of type 2 cytokines^27,28^ to investigate inflammation and remodelling parameters. Together our results highlight mouse strain dependent differences in type 2 and type 17 inflammation that do not seem to alter the development of remodelling but may impact on deposition of specific collagen subtypes.

## Results

### Chronic allergen-induced immune response dynamics differ between C57BL/6 and BALB/c mouse strains

Aspects of allergen-induced airway inflammation have been extensively studied in mouse models, largely in the context of acute Th2-mediated immune responses.^29,30^ Here, using a model of mixed type 2 and type 17 inflammation, we characterised immune cell dynamics in two common in-bred mouse strains, C57BL/6 and BALB/c, strains known to respond differently in models of airway inflammation and hyper-responsiveness.^26,31,32^ Mice were exposed to a multi-allergen cocktail (House Dust Mite, Ragweed and *Aspergillus fumigatus*; DRA) for 4 or 8 weeks (**Figure 1a**) and inflammation assessed. All mice exposed to DRA had a mixed neutrophilic/eosinophilic inflammation, although neutrophilia was not evident in C57BL/6 mice until week 8 (**Figure 1b**). Neutrophils were still present in the BAL 5 days after the last allergen administration (**Figure 1a, b**), even though their numbers were considerably lower compared to eosinophils (**Figure 1b**). Analysis of T lymphocytes and related populations in the lungs, revealed relatively similar proportions of allergen-induced immune cell accumulation between mouse strains, with gradual increases in ICOS^+^ innate lymphoid cells (ILCs) (**Figure 1c**). BALB/c mice generally had a greater ratio of CD4+ to CD8+ T cells compared to C57BL/6 mice reflected in higher total lung CD4+ T cell numbers (**Supplementary figure 1**), but this ratio did not change due to allergen exposure (**Figure 1c**). In addition, innate populations of γδT cells and ICOS^+^ ILCs were more predominant in BALB/c mice compared to C57BL/6 mice with clear increases in cell numbers in BALB/c mice following allergen exposure at either week 4 or 8 (**Figure 1c and supplementary figure 1**). Despite the increased ILC numbers (**Supplementary figure 1**), CD4^+^ T cells and γδT cells appeared to be the major cytokine producing lymphocyte populations in the lungs of all allergic mice. Exposure to DRA resulted in both IL-17A and type 2 inflammatory responses in the lung, with an increased proportion of IL-4 and IL-17A expressing CD4^+^ T cells in both C57BL/6 and BALB/c mice (**Figure 2a**). Interestingly, the numbers of IL-4^+^ CD4^+^ T cells were reduced from week 4 to week 8 in BALB/c mice (**Figure 2b**), also corresponding to a reduction in eosinophils (**Figure 1b**). Nonetheless, there were enhanced numbers of IL-17A^+^ TCRγδ^+^ T cells and IL-17A^+^ CD4^+^ T as well as IL-4^+^ CD4^+^ T cells on week 4 in BALB/c compared to C57BL/6 mice (**Figure 2b**). Of note, TCRγδ^+^ and CD4^+^ T cells contributed equally to the pool of IL-17A^+^ cells in allergic BALB/c mice, whereas CD4^+^ T cells were the main IL-17A^+^ population in C57BL/6 mice (**Figure 2b**).

**Figure 1:**
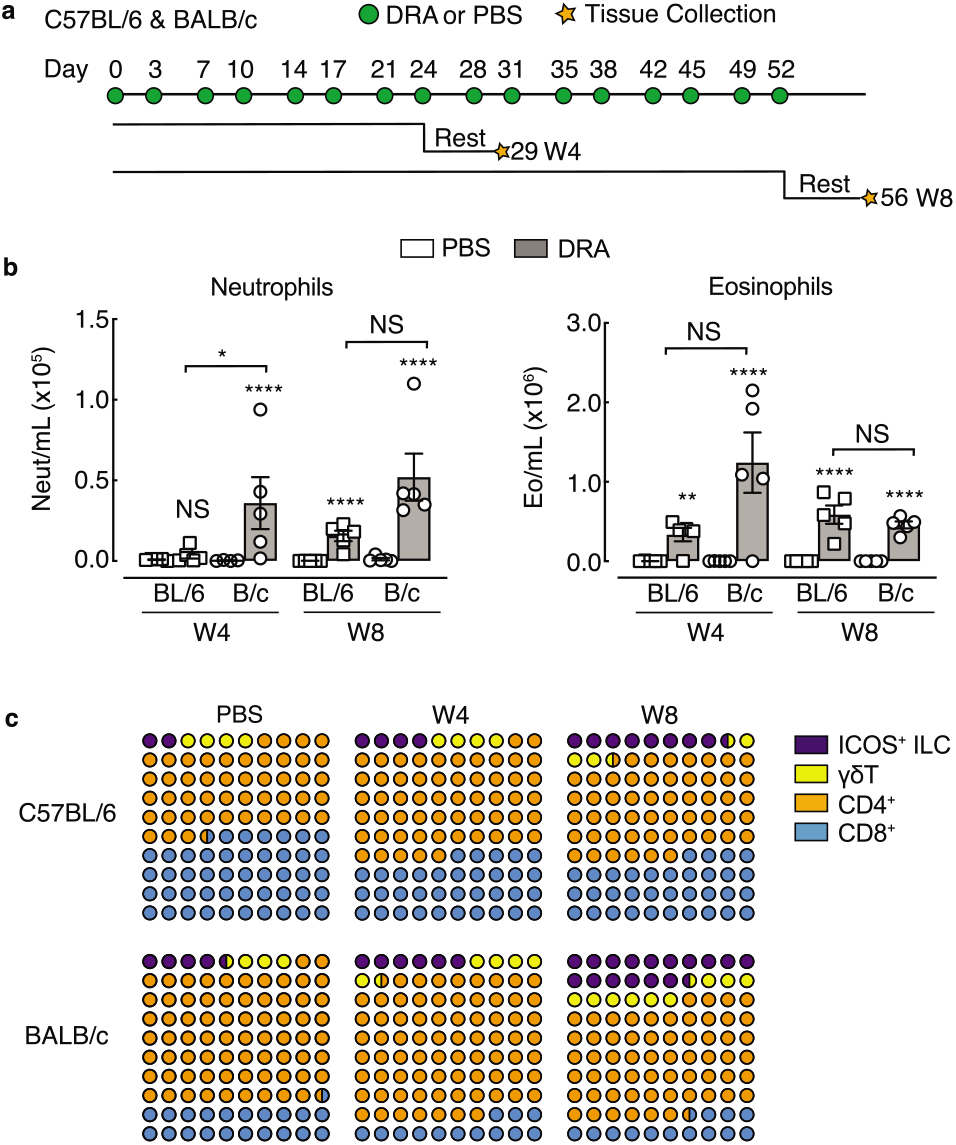
Chronic exposure to DRA allergens induce neutrophil and eosinophil airway inflammation. **a**) Schematic showing allergic airway inflammation model highlighting the timing of DRA or PBS intranasal administration into C57BL/6 or BALB/c mice. Cells were collected for flow cytometry analysis 5 days after the last administration of PBS or DRA (rest) at either 4 or 8 weeks. **b**) Numbers of neutrophils and eosinophils in the BAL of C57BL/6 or BALB/c mice administered PBS or DRA for 4 or 8 weeks. **c**) Plot showing the average proportions of different T cells and ILCs in the lungs of C57BL/6 or BALB/c mice administered PBS or DRA for 4 or 8 weeks. Data are representative of 2 experiments. Data is plotted as mean ± sem with points representing individual animals (**b**). Data in **b** was analysed by ANOVA with Tukey’s multiple comparison test with significance level showing comparisons between either PBS animals within each strain and each time point or C57BL/6 to BALB/c mice as indicated on the graph. NS not significant, \**P*<0.05, \*\*\*\**P*<0.0001.

**Figure 2:**
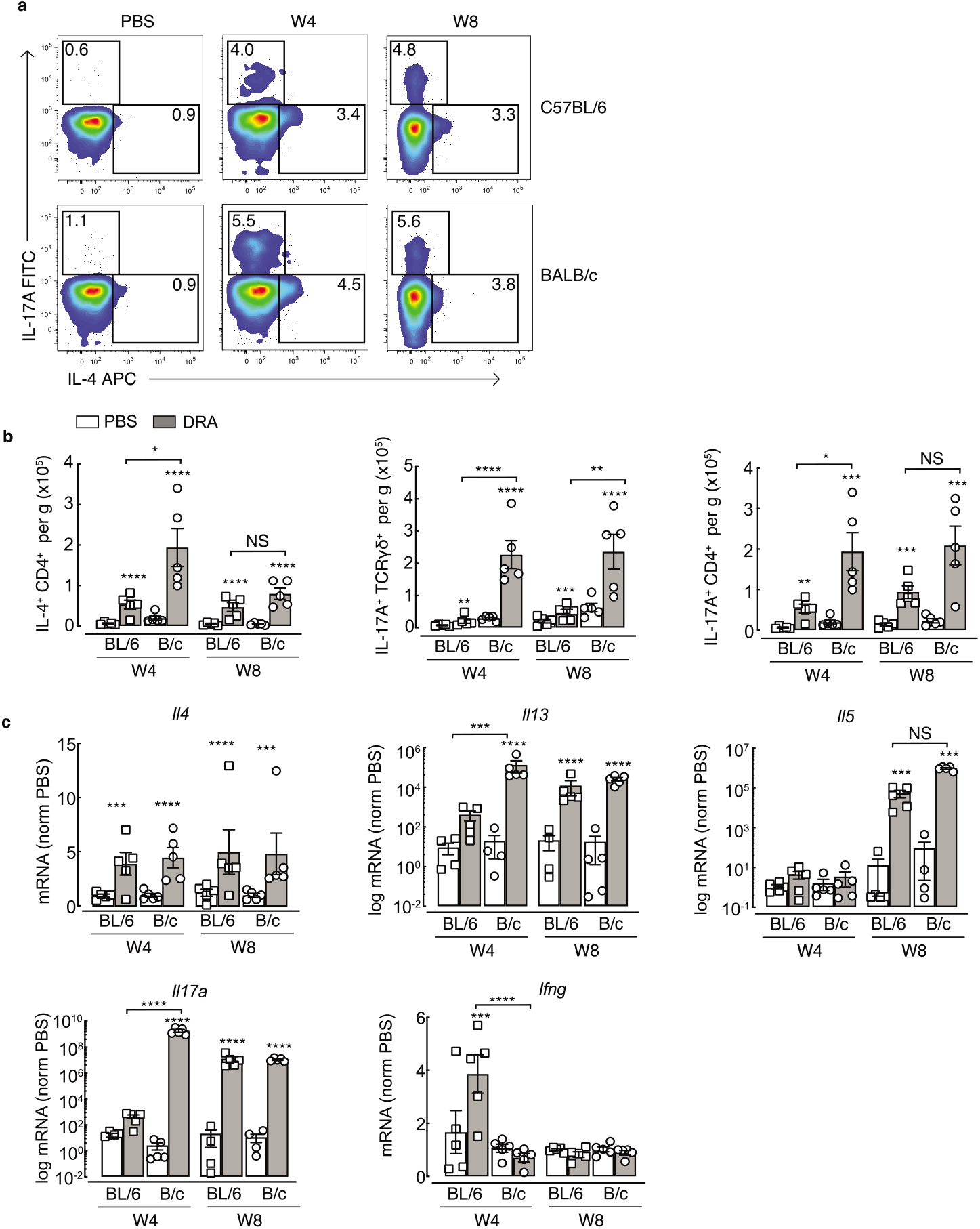
Chronic exposure to DRA allergens induce mixed Th2/Th17 airway inflammation. **a**) Whole lung single cell suspensions were stained for flow cytometry. Live, single TCRγδ^-^TCRβ^+^ CD8^-^CD4^+^ cells were gated on and representative intracellular staining plots of IL-4^+^ and IL-17A^+^ CD4^+^ T cells in the lungs of C57BL/6 or BALB/c mice administered PBS or DRA intranasally two times a week for up to 8 weeks. Cells were analysed by flow cytometry 5 days after the last instillation of allergen. Single cell lung suspensions were stimulated with PMA/ionomycin prior to analysis by flow cytometry. Numbers indicate the percentage of cytokine positive CD4^+^ T cells within each gate. **b**) Absolute numbers of IL-17A^+^ TCRγδ^+^ or IL-17A^+^ or IL-4^+^ CD4^+^ T cells in the lungs of mice as in **a**. **c**) mRNA expression of *Il4, Il13, Il5, Il17a* and *Ifng* in whole lungs of mice treated as in **a.** mRNA were normalised to levels found in PBS C57BL/6 or BALB/c mice at each time point and are relative to geometric mean of housekeeping genes *Gapdh, Rpl13a* and *Rn45s*. Data are representative of 2 experiments. Data is plotted as mean ± sem with points representing individual animals. Data was analysed by ANOVA with Tukey’s multiple comparison test with significance level showing comparisons between either PBS animals within each strain and each time point or C57BL/6 to BALB/c mice as indicated on the graph. NS not significant, \**P*<0.05, \*\**P*<0.01, \*\*\**P*<0.001, \*\*\*\**P*<0.0001.

Expression of key cytokines in whole lung RNA also revealed an increase in type 2 genes (*Il4, Il5, Il13*) and *Il17a* upon DRA treatment with exaggerated *Il13* and *Il17a* expression at week 4 in allergic BALB/c compared to C57BL/6 mice (**Figure 2c**). Whilst we saw no evidence of increased IFNγ^+^ T cells or ILCs (data not shown) in the lungs of mice following allergen administration, there was a transient increase in *Ifng* expression in the lungs of C57BL/6 mice but not BALB/c (**Figure 2c**). Overall, although both mouse strains developed allergen-induced airway inflammation, the degree of inflammation and eosinophilic and neutrophilic responses were higher in BALB/c compared to C57BL/6 mice.

### Chitinase-like proteins are abundantly expressed in the lungs during type 2 and type 17 allergic airway inflammation

Chitinase-like proteins (CLPs) are molecules strongly associated with severe asthma, neutrophilia and IL-17A.^33–36^ Following exposure to DRA allergen, mRNA expression of murine CLPs *Chil1*, *Chil3* and *Chil4* were upregulated in both BALB/c and C57BL/6 mice compared to PBS controls (**Figure 3a**). However, *Chil1* mRNA expression was less in C57BL/6 compared to BALB/c mice, albeit this did not reach statistical significance in this dataset (*P* 0.07 DRA C57BL/6 versus DRA BALB/c mice at week 4 or 8). Additionally, no significant increase in *Chil1* mRNA was detected in whole lung tissue of C57BL/6 mice after 4 weeks of allergen exposure as compared to PBS controls (**Figure 3a**) despite significant increases in secreted BRP-39 protein levels in the BAL (**Figure 3b**). *Chil3* and *Chil4* were significantly increased in both mouse strains at week 4, and expression levels did not change upon further allergen exposure (**Figure 3a**), findings that were supported by measurement of Ym1 secreted protein in the BAL (**Figure 3b**). As there were no commercially available reagents to measure Ym2 protein levels, we developed a Ym2 specific antibody to examine Ym2 expression in the lungs (**Supplementary figure 2**). By western blot, neither Ym1 nor Ym2 was detected in mice administered PBS, but both Ym1 and Ym2 greatly increased following allergen exposure (**Figure 3c**). CLPs can be readily detected in the serum, and serum levels of YKL-40 in humans has been proposed as a biomarker for disease severity and is associated with reduced lung function in several pulmonary pathologies.^37–39^

**Figure 3:**
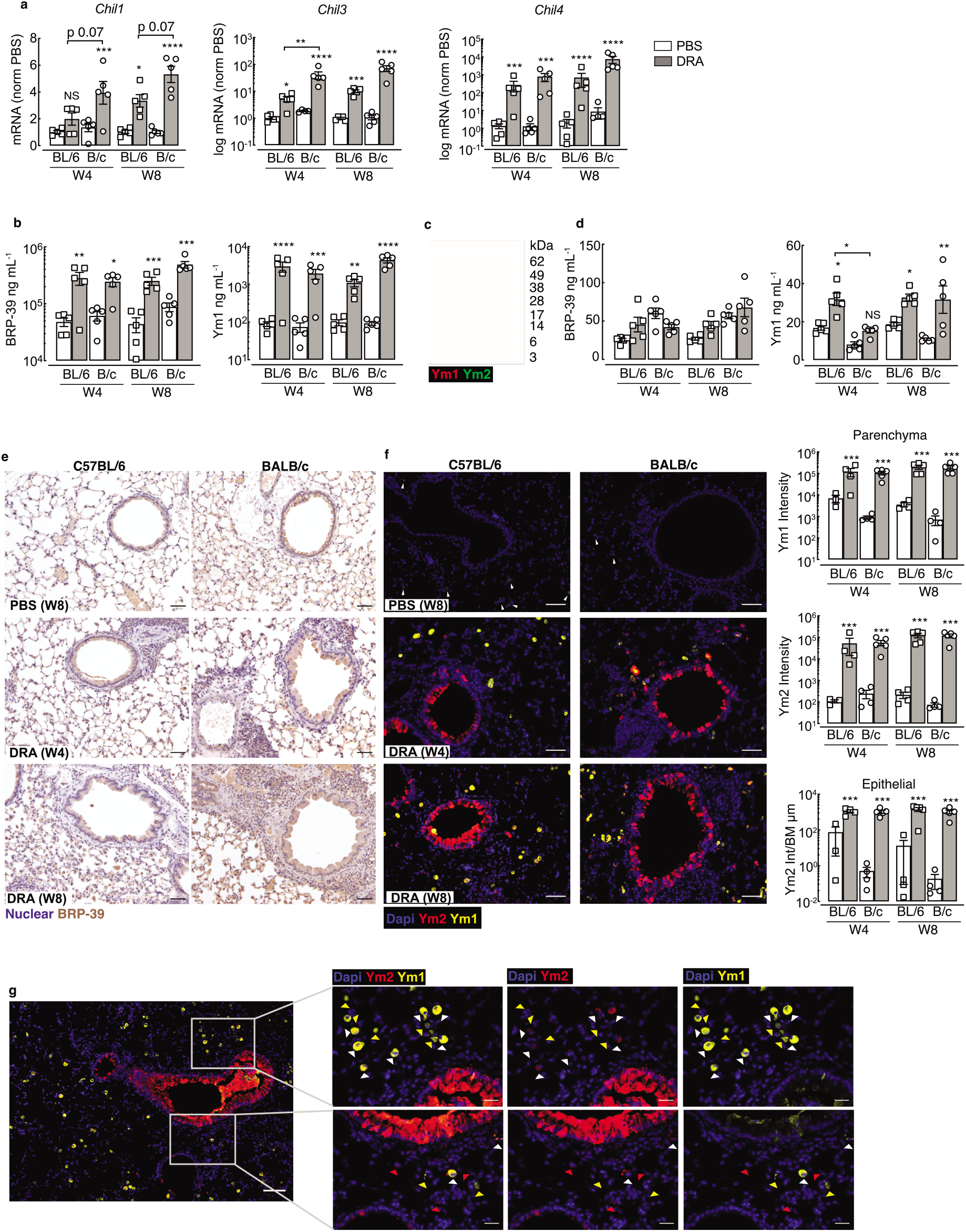
Chitinase-like proteins are abundantly expressed during chronic allergic airway inflammation. **a**) mRNA expression of CLPs *Chil1*, *Chil3* and *Chil4* in whole lungs of C57BL/6 or BALB/c mice exposed intranasally to PBS or DRA for 4 or 8 weeks. Lungs were collected 5 days after the last PBS/DRA administration. mRNA were normalised to levels found in PBS C57BL/6 or BALB/c mice at each time point and are relative to the geometric mean of housekeeping genes *Gapdh, Rpl13a* and *Rn45s. Chil3* and *Chil4* is depicted as log mRNA levels. **b**) Concentration of Ym1 and BRP-39 protein measured by ELISA in the BAL of C57BL/6 or BALB/c mice treated as in **a**. **c**) Western blot analysis of Ym1 (red) and Ym2 (green) levels in the BAL from C57BL/6 mice administered with PBS or DRA for 8 weeks, with BAL taken 5 days after the last DRA/PBS administration. **e**) Concentration of Ym1 and BRP-39 protein measured by ELISA in serum of C57BL/6 or BALB/c mice treated as in **a**. **e**) Microscopy images of immunohistochemical staining of BRP-39 (brown) in lung sections from C57BL/6 and BALB/c mice treated with either PBS for 8 weeks, or DRA for 4 or 8 weeks. Cell nuclei counterstained with haematoxylin (purple); scale bar 50 μm. **f**) Microscopy images of lungs sections of mice as in **a** stained with DNA-binding dye (DAPI) blue; Ym1 (yellow) and Ym2 (red). Scale bar; 50 μm. Images are representative of n=5 mice per group. Quantification of antibody positive staining intensity from stained sections. Ym1 and Ym2 intensity in lung parenchyma areas and Ym2 intensity in airway epithelial cells normalised to length of airway basement membrane. **g**) Microscopy images of immunofluorescent staining for Ym1 (yellow) and Ym2 (red) in lung sections for mice as in **f**. Images show areas where co-staining in airway epithelial or parenchyma cells is evident. Triangles superimposed onto images show Ym1^+^Ym2^-^ cells (yellow), Ym1^-^Ym2^+^ cells (red) or Ym1^+^Ym2^+^ cells (white). Centre image scale bar, 100 μm; outer images scale bar, 50 μm. Datapoints depict individual animals with bars representing mean and sem (**a, b, d, f**). Data are representative of 2 experiments. Data were analysed by ANOVA with Tukey’s multiple comparison test with significance level showing comparisons between either PBS animals within each strain and each time point or C57BL/6 to BALB/c mice as indicated on the graph. NS not significant; *P<0.05, ** P<0.01, ***P<0.001, ****P<0.0001.

Although BRP-39 is a genetic ortholog of YKL-40, the level of BRP-39 in the blood was not significantly altered following allergic-inflammation in this model (**Figure 3d**). However, increased serum Ym1 was detectable in allergic mice of both strains (**Figure 3d**). To determine whether localisation of the three CLPs differed between strains of mice following allergen exposure, we examined immunostained lung sections. BRP-39 was already expressed in macrophages and epithelial cells in the steady state, but the intensity and number of positive cells increased further following allergen administration and the increase was particularly evident in BALB/c mice (**Figure 3e**). Corresponding to secreted levels in the BAL (**Figure 3c**), expression of Ym2 was absent in the lungs of PBS mice, while numerous Ym1^+^ cells, likely alveolar macrophages could be detected (**Figure 3f**).^40^ The expression of Ym1 and Ym2 dramatically increased in the lungs of allergic BALB/c or C57BL/6 mice and the level of expression reached its maximum expression at 4 weeks post DRA treatment (**Figure 3f**). Interestingly, Ym1 and Ym2 appear to have a fairly distinct expression pattern in the lung, with Ym1 largely restricted to myeloid cells and Ym2 largely expressed by epithelial cells, and very few cells that co-stained for Ym1 and Ym2 (**Figure 3f, g**). Overall, we observed modest increases in BRP-39 levels in allergic animals, but strongly enhanced Ym1 and Ym2 expression in the lungs of both allergic C57BL/6 and BALB/c mice. For the first time we show distinct expression of Ym2 in the lungs compared to Ym1, despite their protein sequence being ~96% homologous.

### Allergen-induced immune pathways are fundamentally different between C57/BL6 and BALB/c mouse strains

C57BL/6 and BALB/c mice both developed neutrophilic and eosinophilic airway inflammation in response to chronic allergen administration, despite a greater magnitude of both type 2 and IL-17A cytokine responses in BALB/c mice. Therefore, to more broadly characterise the differences in immune response between mouse strains, we performed differential gene expression analysis of whole lung RNA after 8 weeks of allergen or PBS administration using the NanoString nCounter Myeloid Innate Immunity Panel (NanoString, Amersham, UK). Principal component analysis (PCA) demonstrated a clear separation in gene signatures not only from exposure to DRA versus PBS, Principal Component (PC) 1, but also mouse strain, explained by PC2 (**Figure 4a**). Investigation of the genes that were significantly altered in the DRA model showed that numerous genes were induced (e.g. type 2 effector molecules *Retnla* and *Arg1*) or inhibited (e.g. basement membrane collagens, *Col4a1*, *Col4a1*) to an equivalent degree in both strains (**Supplementary figure 3a**). Hierarchical clustering also separated a considerable number of genes that were regulated in the same way across the strains, but to a much higher degree in one mouse strain over the other (**Figure 4b**) or expression of genes that were fundamentally different between strains (**Supplementary figure 3b**). As predicted from the allergic cytokine responses and alterations in immune cell infiltration into the lung (**Fig 1 & 2**), it was not surprising that type 2 related genes such as *Il13, Fcer2a, Csf2, Ccl2, Ccl11* were more highly upregulated in whole lung tissue from BALB/c compared to C57BL/6 mice (**Figure 4b**). However, interestingly factors known to play an important role in leukocyte adhesion (*Itgb2, Itgb7, Selplg*) were upregulated in allergic C57BL/6 mice but not BALB/c mice (**Figure 4b**), despite an apparent slower rate of inflammatory cell accumulation in C57BL/6 compared to BALB/c mice (**Figure 1b and supplementary figure 1**).

**Figure 4:**
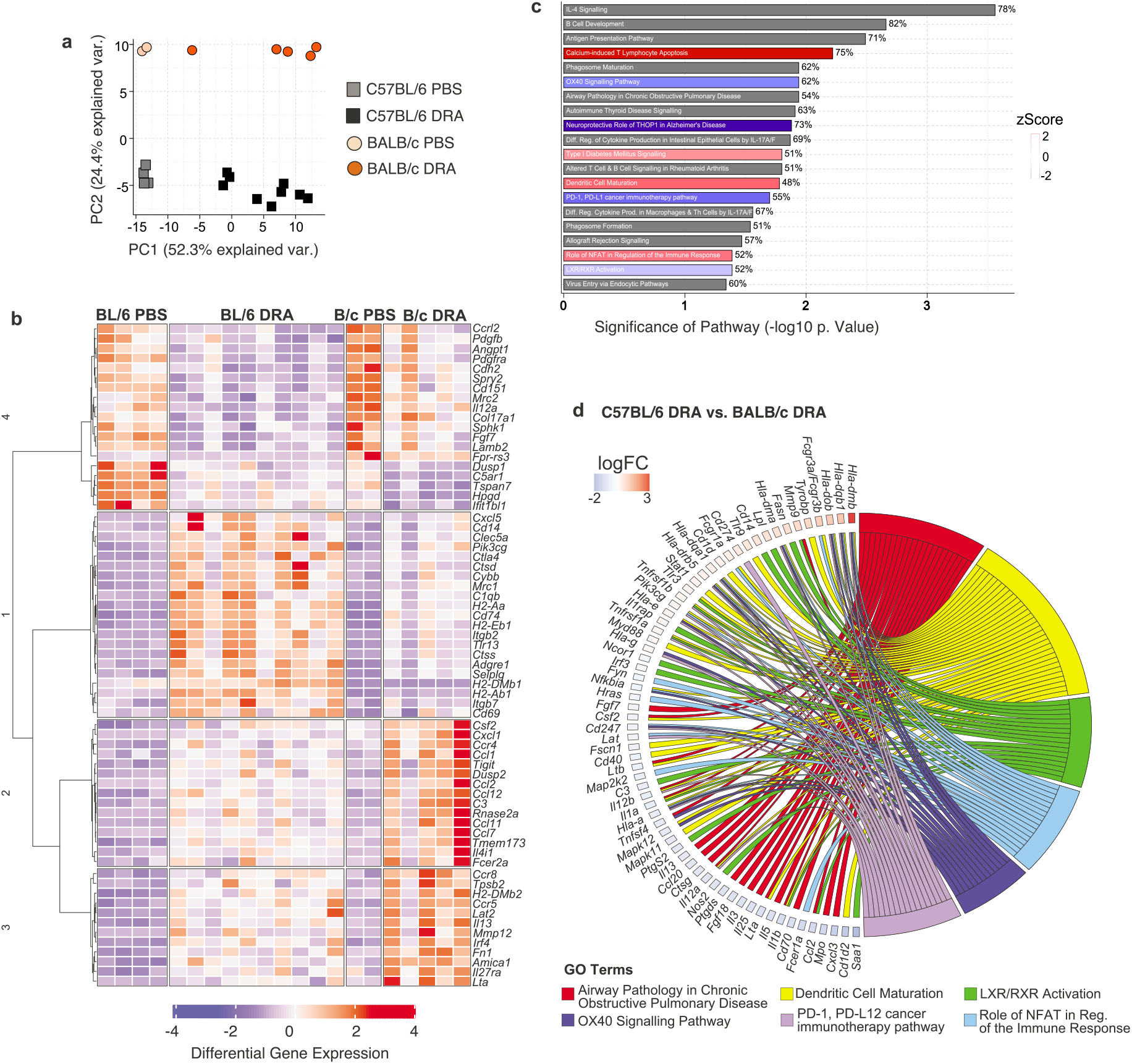
C57BL/6 and BALB/c allergic mice have fundamental differences in immune gene signatures. Whole lung RNA from C57BL/6 and BALB/c mice administered with either PBS or DRA for 8 weeks were analysed using NanoString Myeloid Panel v2. **a**) PCA of expressed genes from C57BL/6 and BALB/c. **b**) Unsupervised, hierarchically clustered heatmap of genes that were significantly regulated in C57BL/6 and BALB/c allergic compared to PBS mice, but also differential regulated between the treated strains. **c**) Differentially expressed genes were visualised with Ingenuity Pathway Analysis tool and top 20 canonical pathways shown for C57BL/6 versus BALB/c mice. Red or blue indicates pathways upregulated or downregulated (respectively) in C57BL/6 compared BALB/c allergic mice. Grey indicates pathways that are significantly regulated but not in a particular direction. Percentage at end of the bar equates to the number of molecules detected compared to the total number of molecules within the canonical pathway. **d**) Chord diagram shows specific genes up or down regulated (colour indicating log fold change) within Go Term that were found to be significantly regulated in C57BL/6 allergic mice compared to BALB/c allergic mice. Transcriptomic analysis was performed on one experiment that was representative of 2 individual experiments.

Analysis of common properties within a signalling pathway (canonical pathway) showed enrichment of various pathways in C57BL/6 compared to BALB/c mice (**Figure 4c**). Pathways including ‘IL-4 signalling’ and ‘airway pathology in COPD’ were significantly different across mouse strains (**Figure 4c**). Whether these pathways were activated or inhibited in C57BL/6 compared to BALB/c mice could not be clearly defined by the analysis (as denoted by the grey bar; **Figure 4c**). However, the specific genes that contributed to the z scores (**Figure 4c**) were also examined (**Figure 4d**). For example, a downregulation of both type 2 cytokines *Il5*, *Il13* and the type 2 inducing cytokine *Il25* in addition to reduced expression of pro-inflammatory cytokines *Il1a, Il1b, Il12a* and *Il12b* indicates that genes characteristic of the “airway pathology in COPD’ pathway (**Figure 4d**) were reduced in allergic C57BL/6 mice relative to BALB/c mice. Interestingly, both ‘OX40 signalling’ and ‘PD-1, PD-L1 signalling’, pathways involved in maintenance and regulation of T cell responses, were downregulated in allergic C57BL/6 compared to BALB/c mice (**Figure 4d**). These and other changes in canonical pathways involved in DC-T cell stimulation possibly explain reduced cytokine production in C57BL/6 mice (**Figure 2**). In addition, LXR/RXR activation, which maintains cholesterol homeostasis but is also known to be anti-fibrotic and anti-inflammatory^41^, was downregulated in C57BL/6 mice (**Figure 4c, d**). Overall, analysis of gene regulation at chronic allergic inflammatory time points revealed differences in gene signatures between mouse strains that may explain reduced immune responses in C57BL/6 compared to BALB/c mice.

### Airway remodelling develops in both C57BL/6 and BALB/c mice despite different allergic inflammation dynamics and immune signatures

The relationship between inflammation and airway remodelling in asthma is still controversial (reviewed by Saglani & Lloyd^42^, Boulet^43^, Guida & Riccio^44^). Some features of remodelling may occur in parallel or even prior to excessive inflammation^18–20^ although difficult to test in the clinical setting. Considering different immune cell dynamics between BALB/c and C57BL/6 mice (**Figure 1–4**), we sought to determine whether features of airway remodelling also varied between mouse strains. Goblet cell hyperplasia is a key feature of remodelling in asthma and contributes to excessive airway mucus secretion.

Equivalent increases in periodic acid schiff (PAS) positive cells, indicative of goblet cells, were observed in both C57BL/6 and BALB/c mice at weeks 4 and 8 (**Figure 5a**). Airway remodelling in asthmatic patients is also characterized by thickening of the basement membrane and deposition of sub-epithelial extracellular matrix proteins. Following DRA allergen exposure, increased collagen deposition, measured by Masson’s Trichrome stain, was also evident around the airways of both mouse strains (**Figure 5b**). Specific immunostaining for components of the ECM (**Figure 6a, c**) previously described to be regulated in asthma^45–48^, supported increases in total airway collagen following allergen exposure (**Figure 5b**). However, fundamental differences in collagen expression between mouse strains were also evident (**Figure 6a, b**).

**Figure 5:**
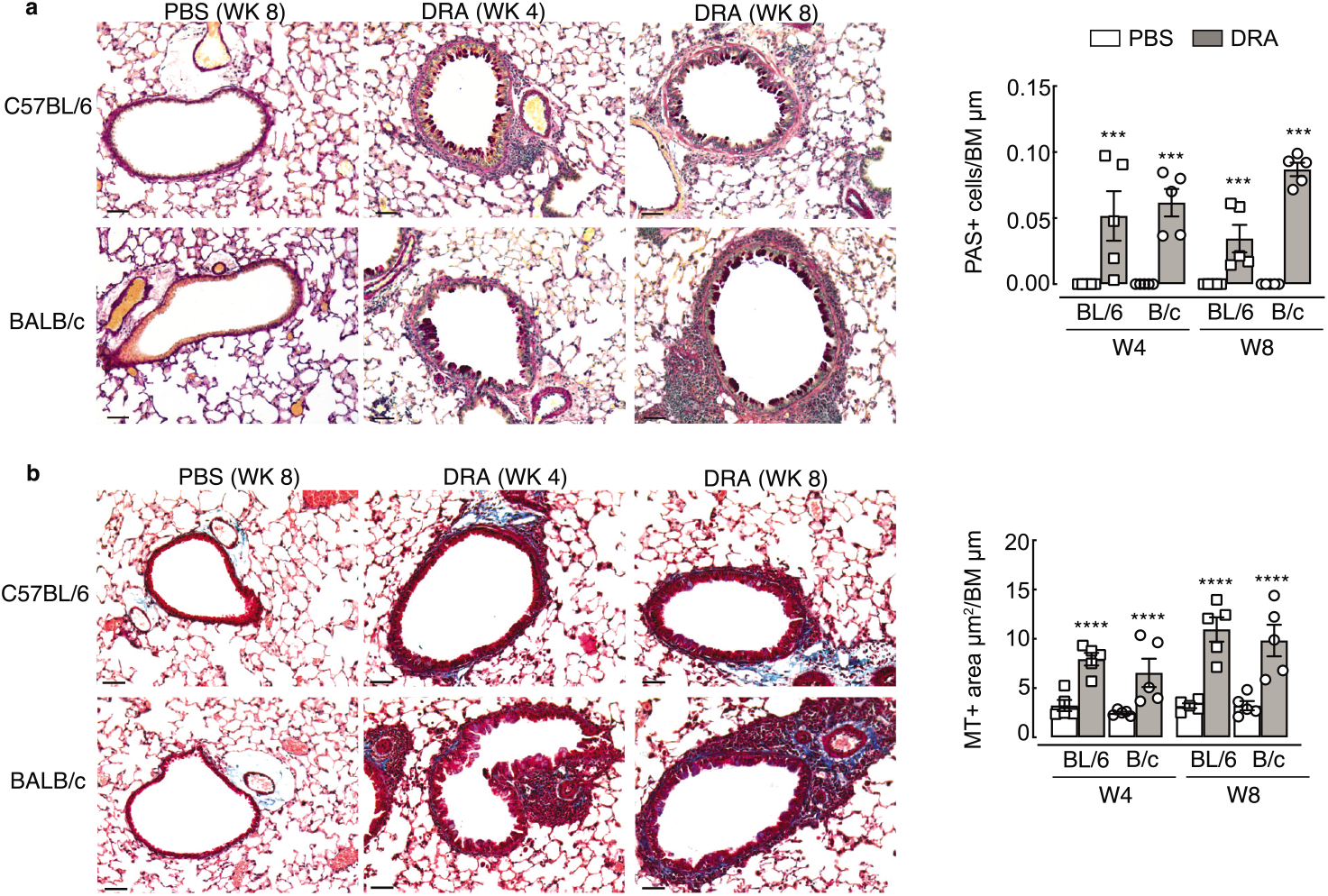
Goblet cell numbers and total collagen increases around the airways following exposure to DRA allergens. C57BL/6 or BALB/c mice were intranasally administered PBS or DRA twice a week up to 8 weeks, and lungs collected for histological analysis 5 days after the last PBS or DRA at weeks 4 and 8. **a**) Microscopy images of lung sections stained for PAS. Airways show PAS^+^ cells (purple) within the epithelium. Graph shows quantification of numbers of PAS^+^ cells per length of basement membrane. **b**) Microscopy images of lung sections stained for Masson’s trichrome from C57BL/6 or BALB/c mice. Airways show accumulation of collagen (blue) below the basement membrane. Graph shows quantification of the area of Masson’s trichrome positive staining around the airways normalised to basement membrane length. All images are representative of n=5 mice; scale bar equals 50 μm. Datapoints depict individual animals with bars representing mean and sem. Data are representative of 2 experiments and were analysed by ANOVA with Tukey’s multiple comparison test and significance level shown relative to PBS animals within each strain and each time point. ***P<0.001, ****P<0.0001.

**Figure 6:**
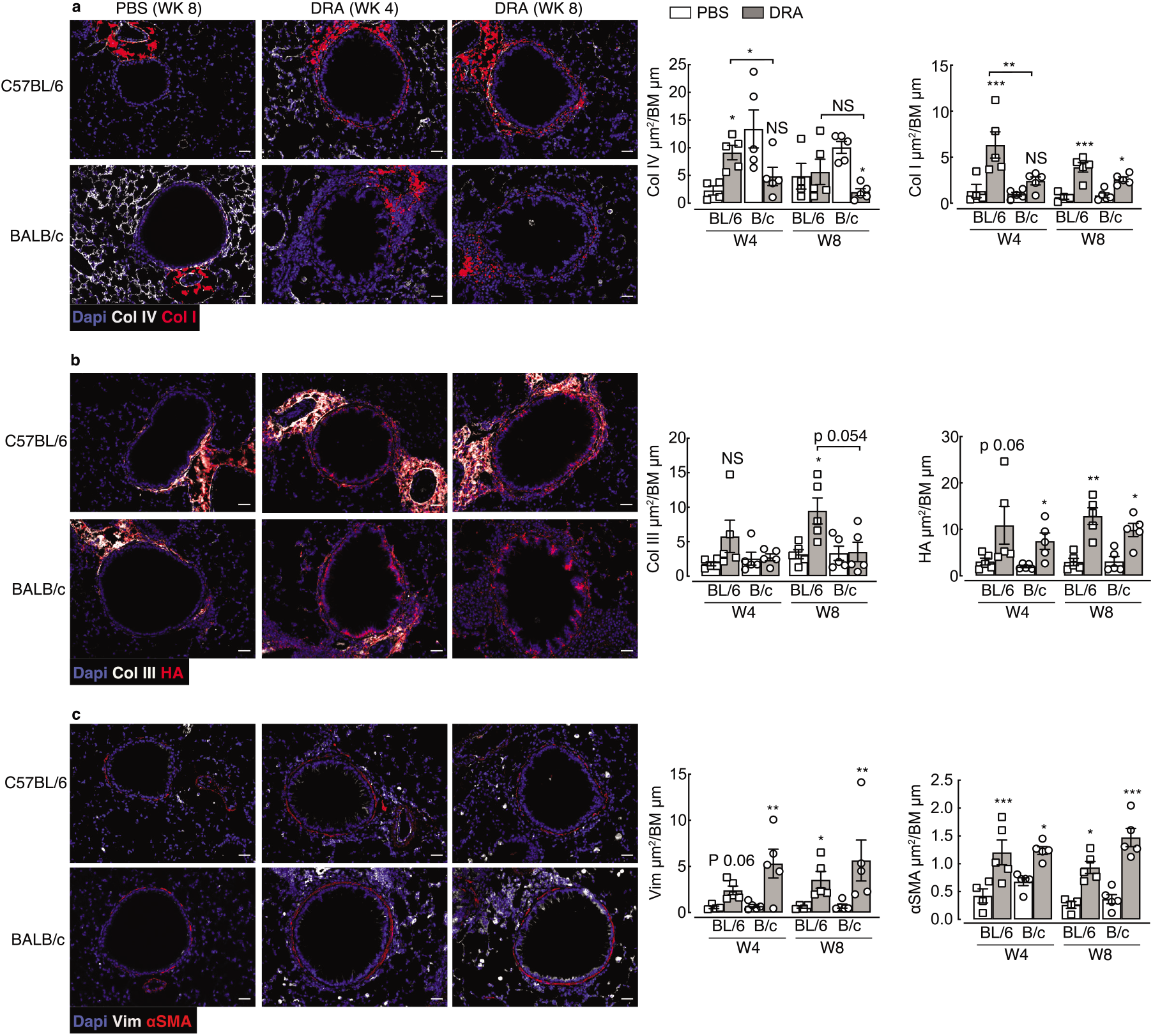
Changes to the ECM and muscle mass around the airway occur following exposure to DRA allergens. C57BL/6 or BALB/c mice were intranasally administered PBS or DRA twice a week up to 8 weeks, and lungs collected for immuno-staining 5 days after the last PBS or DRA at weeks 4 and 8. **a-c**) Microscopy images of lung sections from C57BL/6 or BALB/c mice stained with DNA-binding dye (DAPI) blue; (**a**) Col IV white; Col I red; (**b**) Col III, white; hyaluronan (HA) binding protein, red; (**c**) vimentin (Vim) white; alpha smooth muscle actin (αSMA) red. Scale bar, 30 μm. Images are representative of n=5 mice. Antibody positive staining area was quantified around the airway and normalised to basement membrane length and values are depicted in **a-c**. Datapoints depict individual animals with bars representing mean and sem. Data are representative of 2 experiments and were analysed by ANOVA with Tukey’s multiple comparison test and significance level showing comparisons between either PBS animals within each strain and each time point or C57BL/6 to BALB/c mice as indicated on the graph. \**P*<0.05, ** *P*<0.01, \*\*\**P*<0.001.

Basement membrane protein collagen IV was highly expressed in the steady-state around the airways and alveoli of BALB/c mice compared to C57BL/6 (**Figure 6a**). Upon allergen administration, collagen IV expression decreased over time in BALB/c mice, whereas levels transiently increased in C57BL/6 mice. Additionally, a greater and more rapid increase in airway collagen I in allergic C57BL/6 compared to BALB/c mice was observed (**Figure 6a**) and similarly, accumulation of airway collagen III was significantly increased only in allergic C57BL/6 mice (**Figure 6b**). In contrast, expression of a major glycosaminoglycan component of the ECM, hyaluronan (HA), was increased in response to allergen exposure independently of mouse strain (**Figure 6b**). Changes to collagen composition around the airways of allergic mice was accompanied by an increase in the number of vimentin positive cells (**Figure 6c**), potentially indicating an increase in matrix-secreting fibroblasts.^49^ Additionally, airway muscle mass was also examined and revealed increases in allergic mice regardless of mouse strain (**Figure 6c**). Together, these results demonstrate features of remodelling such as goblet cell hyperplasia, increased smooth muscle mass and ECM changes occur in both C57BL/6 and BALB/c mice. However, differences in deposition of specific collagen subtypes exists between mouse strains.

## Discussion

IL-17 and neutrophilia are often associated with severe asthma.^10^ Despite this, models of allergic airway inflammation still largely focus on studying the regulation of allergen induced type 2 immune responses, utilising BALB/c mice that generally show a strongly skewed type 2 inflammatory response.^26^ Here, we utilised a model of allergic airway inflammation in which neutrophilia and IL-17 are dominant features, with inflammation resistant to steroid intervention ^28^ and AHR unaffected by neutralisation of IL-5 or IL-13 cytokines.^27^ As expected BALB/c mice developed rapid and prominent airway inflammation that was skewed toward type 2 responses, but also greater IL-17 production, particularly by γδ T cells. However, type 2 inflammation was reduced from week 4 to week 8 perhaps reflecting the emergence of a tolerogenic response to allergens in BALB/c mice.^25^ C57BL/6 mice still responded to allergens, but Th2 and IL-17A responses developed at a slower rate compared to BALB/c mice. Delayed type 2 cytokine expression is potentially explained by an early but transient spike in IFNγ expression only observed in C57BL/6 mice. Interestingly, increased IL-17A expression in C57BL/6 mice between weeks 4 and 8 coincided with a reduction in IFNγ levels in allergic mice, which we have shown previously to be an important factor that allows the development of a pulmonary type 2 immune response.^50^ Additionally, IL-10 derived from T cells has been shown to signal via alveolar macrophages leading to suppression of IFNγ-induced airway epithelial disruption.^51^ No difference in expression of *Il10* between strains at chronic time points was observed after DRA administration in our study. However, temporal changes in IL-10 in C57BL/6 mice may contribute to suppression of IFNγ alongside IL-17A.

IL-13 production is known to be higher in BALB/c versus C57BL/6 mice^26^, as also shown here, and is thought to account for increased AHR observed in BALB/c compared to the relatively hypo-responsive C57BL/6 mice.^22,52^ In fact, type 2 cytokine producing cells, rather than eosinophilic inflammation, appear to be key for the maintenance of AHR in models of type 2 airway inflammation.^53,23^ In addition to enhanced type 2 cytokine production, pathway analysis suggested a reduced capacity (in C57BL/6 mice) for antigen-presenting cell (APC) mediated activation of T cells via costimulatory molecules PD-1/PDL-1 and OX40/OX40L, despite enhanced ‘DC maturation’ pathways from NanoString analysis in these mice. Different DC subsets can dictate the allergic immune response and targeting either DC activation or molecules involved in antigen presentation may be a fruitful approach to therapeutically target allergic asthma, and specifically different immune phenotypes of disease.^54,55^ PDL1 is known to enhance AHR and Th2 cytokine production in allergic mice.^56^ Thereby a reduction in PD1-PDL1 signalling, alongside reduced OX40 signalling may explain the reduced Th2 response in C57BL/6 mice compared to BALB/c. Future research identifying specific dendritic cell phenotypes in both strains during AAI could prove useful for understanding pathways to target AHR and inflammation in asthma.

In this model of Th2/Th17 allergic airway inflammation, both BALB/c and C57BL/6 mice developed a similar degree of airway remodelling in response to allergen exposure. However, our study reveals intriguing differences in ECM composition between mouse strains, not only in response to allergens but also in the steady state. One could anticipate that changes in collagen composition, particularly centred around the ratios of collagen I and III, could profoundly alter lung function and along with varied immune responses, may contribute to well reported differences in AHR measurement between mouse strains.^22,52^ Both collagen I and III play major roles in the structural integrity of tissues and are often co-expressed within the tissue, with Col I contributing to tensile strength, whilst Col III allows tissue flexibility.^57^ Collagen III can modulate scar formation^58^ and during early active fibrosis levels of Col III significantly increase.^59,60^ However, an increased ratio between type I and type III collagen occurs in infants diagnosed with chronic lung disease proceeding respiratory distress syndrome.^59^ Additionally, a lack of collagen III can disturb the development of collagen fibril formation resulting in functional failure of the organ.^61,62^ Here, allergic BALB/c mice appeared to have preferential increase in collagen I around the airways, with no significant changes to collagen III, although we cannot rule out expression of collagen III at time points earlier than week 4. A failure to induce collagen III during remodelling processes may in fact perturb lung function, perhaps contributing to increases in AHR often observed in BALB/c mice in response to allergen challenge.^22,26,52^ Differential dynamics in col IV expression between mouse strains is also intriguing, as Col IV is crucial for barrier formation anchoring airway epithelial cells. The rapid loss of Col IV in BALB/c mice may relate to a significant increased vimentin-positive cells around the airway at week 4, and potentially enhanced epithelial mesenchymal transition leading to a more rapid remodelling response in BALB/c versus C57BL/6. Although both mouse strains feature a similar magnitude of allergen-induced remodelling, further analysis of the early dynamics, pre-week 4, and mechanisms leading to changes in the ECM in these two mouse strains may reveal important features of tissue remodelling in disease.

Remodelling is typically examined as a change in epithelial goblet hyperplasia, increased muscle mass and total collagen, but here our study in genetically distinct mouse strains, highlights that the term remodelling is much more complicated. Just as inflammation varies greatly between asthmatic cohorts, airway remodelling too may be considered an “umbrella” term, whereby different pathways are likely to be more or less important in different asthma phenotypes. A greater understanding of how ECM composition changes can alter lung mechanics/function, but also how the differing ECM components can regulate immune cell recruitment and activation, will help us to understand the development of lung diseases like asthma and whether approaches to target remodelling will prove useful in treating such chronic inflammatory diseases. Furthermore, it is interesting to speculate that different genetic strains of mice, rather than using different allergens of timings of allergen exposure, could prove more useful for modelling different trajectories of allergic asthma in people.

## Methods

### Animals and Ethics

Wild-type (BALB/c or C57BL?6J) mice were obtained from a commercial supplier (Envigo, Hillcrest, UK). Experimental mice, all female, were between 7-10 weeks old at the start of the experiment and were housed in individually ventilated cages maintained in groups of 5 animals in specific pathogen-free facilities at the University of Manchester. Mice were not randomised in cages, but each cage was randomly assigned to a treatment group. Sample size was calculated on the basis of the number of animals needed for detection of a 25% change in Masson’s trichrome positive area around the airway in PBS versus allergic mice, with a P value of <0.05, based on pilot experiments carried out with 3 mice per group. All animal experiments were performed in accordance with the UK Animals (Scientific Procedures) Act of 1986 under a Project License (70/8548) granted by the UK Home Office and approved by the University of Manchester Animal Welfare and Ethical Review Body. Euthanasia was performed by asphyxiation in a rising concentration of carbon dioxide.

### Model of allergic airway inflammation

Allergic airway inflammation was induced in mice in a similar manner as has been described previously.^27^ Allergen DRA cocktail comprising of 5 μg House Dust Mite (*Dermatophagoides pteronyssinus*, 5450 EU, 69.23 mg per vial), 50 μg Ragweed (*Ambrosia artemisifolia*), 5 μg *Aspergillus fumigatus* extracts (Greer Laboratories, Lenoir, NC, USA) were freshly prepared prior to each inoculation. Mice were briefly anaesthetised via inhalation of isoflurane, and 20 μL of DRA cocktail or PBS were given via intranasal inoculation twice weekly for up to 8 weeks. Mice were rested for 5 days prior to performing BAL and collecting lung tissue.

### Isolation of cells from the BAL and lung tissue

Following exsanguination, BAL cells were obtained through cannulation of the trachea and washing the lungs with 0.4 mL PBS (Sigma Aldrich, St. Louis, MO, USA) containing 0.25 % BSA (Sigma Aldrich) (four washes). Lungs were processed as previously described.^40^ Briefly, a right lobe was removed and minced in 1 mL of HBSS buffer containing 0.4 U mL^-1^ Liberase TL (Sigma Aldrich) and 80 U mL^-1^ DNase type I (ThermoFisher Scientific, Waltham, MA, USA) for 25 min in a 37°C shaking incubator. Digestion was stopped with 2 % FBS (ThermoFisher Scientific) and 2 mM EDTA prior to passing the suspension through a 70 μm cell strainer (Greiner Bio-One, Stonehouse, UK). Red blood cells were lysed (Sigma) and total live BAL and lung cell counts assessed with Viastain AOPI (Nexcelom Bioscience LLC, Lawrence, MA, USA) using a Cellometer Auto2000 automated cell counter (Nexcelom Bioscience LLC).

### Flow Cytometry

Equal cell numbers of each lung and BAL sample were stained for flow cytometry. Cells were washed with ice-cold PBS and stained with Live/Dead Aqua or Blue (ThermoFisher Scientific) for 10 min at room temperature. All samples were then incubated with Fc block (5 μg mL^-1^ CD16/CD32 (BD Biosciences, San Diego, CA, USA) and 0.1 % mouse serum in FACs buffer (PBS containing 0.5 % BSA and 2 mM EDTA (ThermoFisher Scientific)) for 20 min before staining for specific surface markers with fluorescence-conjugated antibodies for 25 min at 4°C (**Table 1**) Following surface staining, cells were fixed with ICC fix (Biolegend, San Diego, CA, USA) and stored at 4°C until intracellular staining was performed or cells were acquired. For intracellular cytokine staining, cells were stimulated for 4h at 37°C with PMA (phorbol myristate acetate; 0.5 μg mL^-1^; Sigma Aldrich) and ionomycin (1 μg mL^-1^)(Sigma Aldrich) and for 3 h at 37°C with Brefaldin A (10 μg mL^-1^; Biolegend). Cell surfaces were stained and cells fixed as described above. All cells were permeabilized (eBioscience, San Diego, CA, USA) then stained with antibodies for intracellular cytokines (**Table 1**). Cells were identified with the following markers: eosinophils F4/80^+^ CD11c^-^ CD11b^+^ SigF^+^; neutrophils Ly6G^+^ CD11b^+^ CD11c^-^; T cells TCRβ^+^ TCRγδ^-^ and either CD4^+^ or CD8^+^; gamma delta T cells TCRβ^-^ TCRγδ^+^ CD4^-^CD8^-^; innate lymphoid cells (ILCs) CD90^+^ ICOS^+^ Lineage^-^ (CD11b, Ly6G, Ly6C, CD11c, Ter119, NK1.1, B220, CD3). All samples were acquired with a FACS Canto II or 5 laser Fortessa with BD FACS Diva software and analysed with FlowJo software (versions 9 and 10; BD Biosciences).

**Table 1:** Antibodies used for flow cytometry analysis

| Antigen | Antibody Clone | Isotype | Source |
| --- | --- | --- | --- |
| *Ly6G* | 1A8 | Rat IgG2a κ | Biolegend |
| *CD11b* | M1/70 | Rat IgG2b κ | Biolegend |
| *CD11c* | N418 | Armenian Hamster IgG | Biolegend |
| *F4/80* | BM8 | Rat IgG2a κ | Biolegend |
| *CD64* | X54-5/7.1 | Mouse IgG1 κ | Biolegend |
| *SiglecF* | E50-2440 | Rat IgG2a κ | BD Biosciences |
| *I-A/I-E* | M5/144.15.2 | Rat IgG2b κ | Biolegend |
| *TCRβ* | H57-597 | Armenian Hamster IgG | eBioscience |
| *TCRγδ* | GL3 | Armenian Hamster IgG | Biolegend |
| *CD4* | GK1.5 | Rat IgG2b κ | Biolegend |
| *CD8* | 53-6.7 | Rat IgG2a κ | Biolegend |
| *CD3* | 17A2 | Rat IgG2b κ | Biolegend |
| *B220* | RA3-682 | Rat IgG2a κ | Biolegend |
| *Ter119* | Ter-119 | Rat IgG2b κ | Biolegend |
| *NK1.1* | PK136 | Mouse IgG2a κ | Biolegend |
| *ICOS* | C398.4A | Armenian Hamster IgG | Biolegend |
| *CD90.2* | 30-H12 | Rat IgG2b κ | Biolegend |
| *IL-4* | 11b11 | Rat IgG1 κ | Biolegend |
| *IL-13* | eBio13A | Rat IgG1 κ | eBioscience |
| *IL-17a* | TC11-18H10.1 | Rat IgG1 κ | Biolegend |
| *IFNγ* | XMG1.2 | Rat IgG1 κ | Biolegend |

### RNA extraction and qRT-PCR

One right lung lobe was stored in RNAlater (ThermoFisher Scientific) prior to homogenization in Qiazol reagent (Qiagen, Hilden, Germany). RNA was prepared according to manufacturer’s instructions and stored at −70 °C. Reverse transcription of 0.2-0.5 μg total RNA was performed using 50 U Tetro reverse transcriptase (Bioline, London, UK), 40 mM dNTPs (Promega), 0.5 μg primer for cDNA synthesis (Sigma Aldrich) and RNasin inhibitor (Promega, Madison, WI, USA). The transcripts for genes of interest were measured by real-time PCR with a Lightcycler 480 II system (Roche, Basel, Switzerland) and a Brilliant III SYBR Green Master mix (Agilent Technologies, Santa Clara, CA, USA) with specific primer pairs (**Table 2**). mRNA amplification was analysed by second derivative maximum algorithm (LightCycler 480 Sw 1.5; Roche) and expression of the gene of interest was normalised to the geometric mean of three housekeeping genes *Rn45s, Rpl13a, Gapdh* (**Table 2**).

**Table 2:**
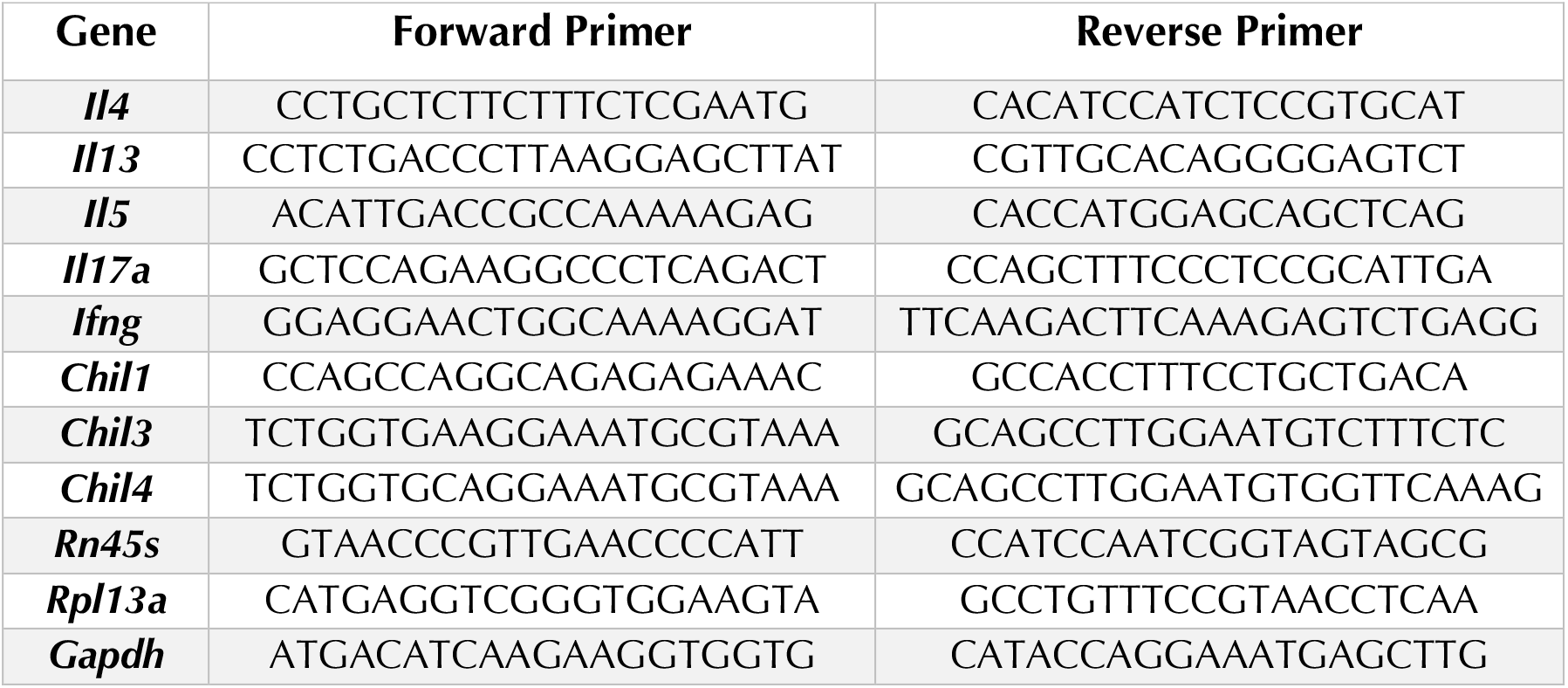
Sequences of primers for measurement of mRNA expression via quantitative RT-PCR

### Transcriptome profile and associated analysis

Quality of RNA extracted from lung tissue, as described above, was assessed with Agilent 2200 TapeStation system prior to downstream analyses, samples with an RIN value of <5.5 were excluded. RNA concentration was determined using Qubit TM RNA BR Assay Kit (ThermoFisher Scientific) and 100 ng RNA (per sample) run on a Nanotstring nCounter R FLEC system using the Myeloid Innate Immunity v2 panel (XT-CSO-MMII2-12). Note, the probes in this panel do not distinguish between *Chil3* and *Chil4*. Raw counts were uploaded onto nSolver version 4.0 using default settings. Non-normalised counts were exported, and subsequent analyses performed in R (version 3.6.3) using RStudio Version 1.2.5033 (2009-2019 RStudio, Inc, Boston, MA, USA). Positive controls were analysed to ensure there was clear resolution at variable expression levels and negative controls were used to set a minimum detection threshold which was then applied to all samples. Data were normalised with EdgeR using the Upper Quartile method and differential expression of genes calculated via linear modelling accounting for sample quality weights with Empirical Bayes smoothing using the limma-voom R packages.^63^ All genes expressed above the background threshold were used for principal component analysis (PCA). Genes with an absolute fold change of greater than 0.5 and a significance value of under 0.05 after correction for multiple comparisons using the Benjamini-Yekeuteli method were defined as “differentially expressed” and taken forward for further analysis. Heatmaps were then generated from scaled normalized counts of DE genes using the ComplexHeatmaps R package. The networks and functional analyses of DE genes were generated with Ingenuity Pathway Analyser (IPA; QIAGEN Inc., https://www.qiagenbio-informatics.com/products/ingenuity-pathway-analysis). Within the IPA software no tissue filtering was used and the user dataset was defined as the reference. Pathway data were then imported into R for visualisation using the ggplot package.

### Generating Anti-Ym2 and determining antibody specificity

Anti-Ym2 specific antibodies were generated by Cambridge Research Biochemicals (Billingham, UK). The 9 amino acid sequence at the N-terminal (CKASYRGEL) were used as the immunogen as it has almost no homology to the Ym1 sequence. Bacterial optimised expression plasmids for Ym1 and Ym2 were purchased from (Genscript, Piscataway, NJ, USA). Plasmids were then transfected into competent *E. Coli* (BL21) using heat shock followed antibiotic selection against Ampicillin (Amp; 25 mg mL^-1^) and Chloramphenicol (Chl; 34 mg mL^-1^). To generate recombinant protein a small scraping of the stock sample was expanded in LB media including antibiotics until optical density reached between 0.6-1.0 at which point IPTG (0.1 M) was added to the cultures. The OD was kept under 1.0 by diluting the culture with fresh media as required and left overnight. Thereafter bacteria were pelleted and resuspended in loading buffer containing DTT (200 mM; ThermoFischer Scientific).

### Western Blotting

Lysed Ym1 or Ym2 transfected *E.coli* cells and murine BAL were denatured in the presence of DTT (200 mM; Thermo Fischer Scientific) for 5 mins at 95°C. Each sample (2-10 μL) or protein ladder (Seeblue; ThermoFischer Scientific) was separated on Bis-Tris 4-12 % gradient gel with MES buffer (ThermoFischer Scientific) before transfer onto a PVDF membrane. The membrane was washed in distilled water followed by incubation in blocking buffer (5 % BSA in PBST (0.05 % Tween-20 in PBS)) for 60 mins at room temperature on a rocking platform. Primary antibodies were used at 1:500 (rabbit anti-mouse Ym2, polyclonal (custom made) or goat anti-mouse Ym1 polyclonal; R&D Systems, Minneapolis, MN, USA) and incubated at room temperature over-night on a rocking platform. The membrane was then washed in PBST followed by secondary antibody detection (1:1000 anti-Rabbit IgG Cy3 and Streptavidin-Cy3; ThermoFisher Scientific) for one hour at room temperature. Membranes were imaging using a Gel Doc (Azure Biosystems, Cambridge Bioscience, Cambridge, UK).

### Histology and Immunostaining

The left lung lobe was fixed perfused with 10 % neutral buffered formalin (Sigma Aldrich) and was incubated overnight before being transferred to 70 % ethanol. Lungs were processed and embedded in paraffin, then sectioned (5 μm) and stained with Masson’s trichrome (MT) or periodic acid schiff (PAS) stains using standard protocols. Images were captured with a Leica microscope with digital DMC2900 camera. For immunostaining with antibodies, lung sections were deparaffinised and heat-mediated antigen retrieval performed using Tris-EDTA buffer (10 mM Tris Base, 1 mM EDTA, 0.05 % Tween-20 pH 8.0; incubation 20 min 95°C). Non-specific protein was blocked with 2 % normal donkey serum (Sigma Aldrich) in PBS containing 0.05 % Tween-20 and 1 % BSA. If a biotin labelled antibody or probe was used, avidin biotin block (ThermoFisher Scientific) was performed prior to an overnight incubation at 4°C with primary antibodies (Table 3). Sections were washed in PBS before incubation with secondary antibodies (**Table 3**) for 1 hr at room temperature followed by mounting with DAPI containing fluoromount (Southern Biotech, Birmingham, AL, USA). Images were captured with an EVOS FL imaging system (ThermoFisher Scientific). Analysis of images was performed using ImageJ software (version 2.09.0-rc69/1.52p) on sections where sample identification was blinded for the investigator and airways analysed had to be intact and fit within a single microscope field of view 480 μm x 360 μm. Goblet cells were visualised on PAS-stained sections and numbers of PAS+ cells counted per airway and normalised to the length the airway basement membrane. Total collagen area was calculated by measuring the area of Masson’s trichrome positive stain (blue) around the airway and values were normalised to basement membrane length. For calculation of collagen, hyaluronan, vimentin and αSMA area, background autofluorescence was subtracted from all images based on pixel intensities of sections stained with secondary antibodies only. A region of interest was drawn parallel to the airway basement membrane at a distance of 50 μm. A threshold was applied to all images to incorporate positively stained pixels and area of positively stained pixels within the region of interest calculated and normalised to the length of the basement membrane. All areas of the airway that contained a blood vessel was excluded from analysis to ensure measurements specifically related to airways and not vasculature. For all image analysis, between 5-15 airways were measured per mouse.

**Table 3:** Antibodies used for immuno-histological analysis

| **Antigen** | **Antibody Clone** | **Dilution** | **Source** |
| --- | --- | --- | --- |
| ***Ym1*** | Goat polyclonal - Biotinylated | 1:100 | R&D – BAF2446 |
| ***Ym2*** | Rabbit polyclonal | 1:1000 | Home-made |
| ***Collagen I*** | Goat polyclonal | 1:200 | Cambridge Bioscience – 1310-01 |
| ***Collagen III (N-terminal)*** | Rabbit polyclonal | 1:300 | Proteintech – 22734-1-AP |
| ***Collagen IV alpha I*** | Rabbit polyclonal | 1:200 | Novus Biologicals – NB120-6586 |
| ***Hyaluronan binding protein*** | Biotinylated | 1:100 | Merck Millipore - 385911 |
| ***α-Smooth muscle actin*** | Goat polyclonal | 1:200 | Novus Biologicals – NB-300-978 |
| ***Vimentin*** | Rabbit polyclonal | 1:200 | Abcam – ab45939 |
| ***BRP-39*** | Rabbit Polyclonal | 1:100 | Biorbyt – orb10365 |
| - | Streptavidin 557 | 1:800 | R&D – NL999 |
| - | Streptavidin 637 | 1:400 | R&D – NL998 |
| - | Donkey anti-rabbit IgG 557 | 1:200 | R&D – NL004 |
| - | Donkey anti-rabbit IgG 637 | 1:200 | R&D – NL005 |
| - | Donkey anti-goat IgG 557 | 1:200 | R&D – NL001 |
| - | Donkey anti-goat IgG 637 | 1:200 | R&D – NL002 |

### Quantification of Ym1 and BRP-39

The levels of Ym1 and BRP-39 in the serum and BAL were measured by sandwich ELISA, with DuoSet ELISA kits (R&D Systems) as per manufacturers recommendation.

### Statistical analysis

Statistical analysis was performed using JMP Pro 12.2.0 for Mac OS X (SAS Institute Inc., Cary, NC, USA). Normal distribution of data was determined by optical examination of residuals, and each group was tested for unequal variances using Welch’s test. Differences between groups were determined by analysis of variance (ANOVA) followed by a Tukey-Kramer HSD multiple comparison test or unpaired two-tailed Student’s t-test as indicated in figure legends. In some data sets, data were log transformed to achieve normal distribution. Differences were considered statistically significant for *P* values of less than 0.05.

## Acknowledgments

This work was supported by the Medical Research Foundation UK jointly with Asthma UK (MRFAUK-2015-302 to TES), the Medical Research Council UK (MR/P02615X/1 to DR and MR/K01207X/1 to JEA) and the Wellcome Trust (106898/A/15/Z to JEA). The authors would like to thank Brian Chan for his technical support, Alistair Chenery and Anthony Day for *E. coli* expressing recombinant Ym1 and Ym2 and Conor Finlay for critical reading of the manuscript. We also thank the Flow Cytometry, Histology and Biological Services core facilities at the University of Manchester.

## Author Contributions

**JEP**: Data curation; investigation; methodology; formal analysis; writing – original draft; Writing – reviewing and editing. **SP**: investigation; methodology. **DR:** investigation; writing – reviewing and editing. **JEA**: Funding acquisition; investigation; writing – reviewing and editing. **TES**: Conceptualisation; Investigation; methodology; project administration; supervision; funding acquisition; writing – original draft; writing – review and editing.

