## Supplementary Figures for "The magnitude of airway remodelling is not altered by distinct allergic inflammatory responses in BALB/c vs C57BL/6 mice but matrix composition differs"

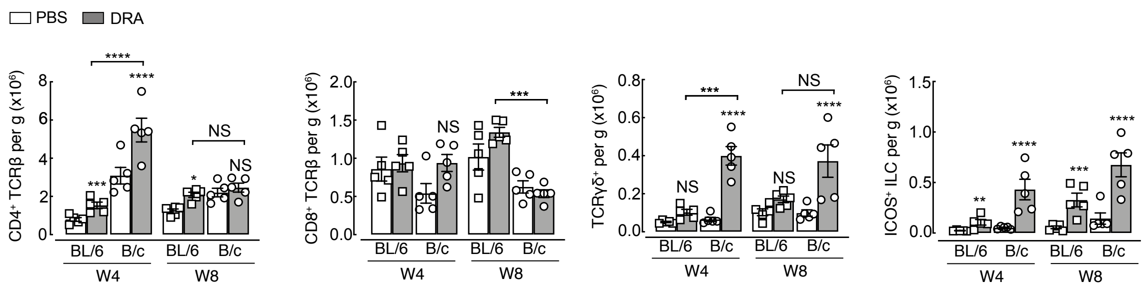


**Supplementary figure 1:** C57BL/6 or BALB/c mice were intranasally administered PBS or DRA twice a week up to 8 weeks, and cells collected for flow cytometry analysis 5 days after the last administration of PBS or DRA. Numbers per gram of tissue of different T cell populations or ICOS$^{+}$ ILCs in the lungs of C57BL/6 or BALB/c mice administered PBS or DRA for 4 or 8 weeks. Data are representative of 2 experiments. Data is plotted as mean ± sem with points representing individual animals. Data was analysed by ANOVA with Tukey's multiple comparison test with significance level showing comparisons between either PBS animals within each strain and each time point or C57BL/6 to BALB/c mice as indicated on the graph. NS not significant, *$P$<0.05, *** $P$<0.001 and ****$P$<0.0001.

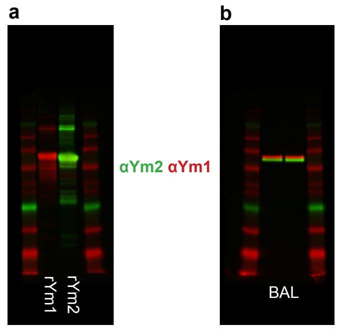


**Supplementary figure 2:** Specificity of anti-Ym1 and anti-Ym2 was analysed by western blot with **a**) total lysates expressing recombinant Ym1 and Ym2 or (**b**) BAL from C57BL/6 mice administered DRA allergens for 8 weeks. Ym1 and Ym2 protein was detected at ~48kDa.

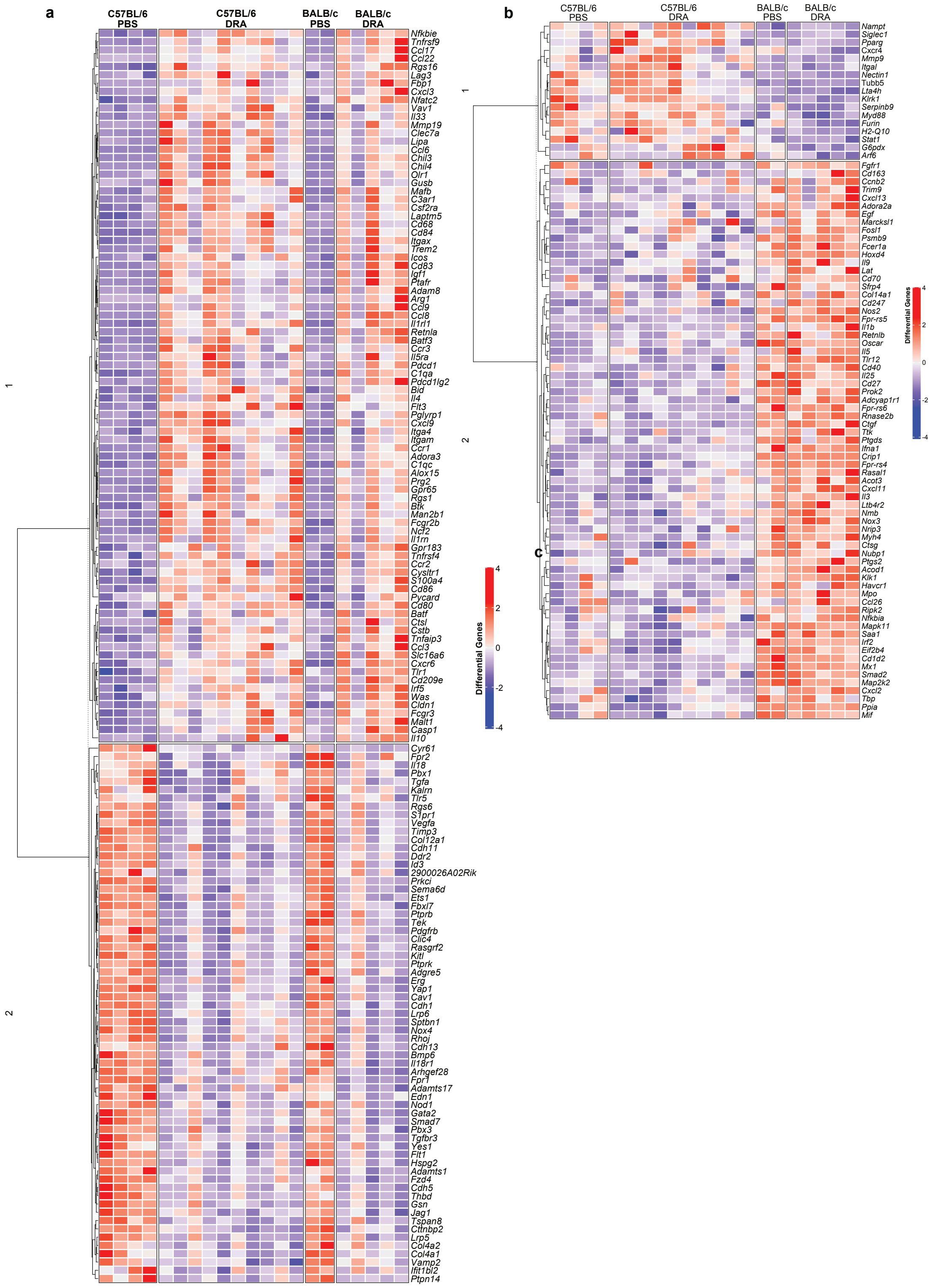


**Supplementary figure 3:** Genes regulated in response to allergic inflammation in C57BL/6 and BALB/c mice. Whole lung RNA from C57BL/6 and BALB/c mice administered with either PBS or DRA for 8 weeks were analysed using NanoString Myeloid Panel v2. Unsupervised, hierarchically clustered heatmap of **a**) differentially expressed genes between PBS and DRA depicted for both mouse strains or **b**) genes that are differentially expressed between strains independently of allergen administration.
